# On the stability and layered organization of protein-DNA condensates

**DOI:** 10.1101/2021.08.22.457249

**Authors:** Andrew P. Latham, Bin Zhang

## Abstract

Multi-component phase separation is emerging as a key mechanism for the formation of biological condensates that play essential roles in signal sensing and transcriptional regulation. The molecular factors that dictate these condensates’ stability and spatial organization are not fully understood, and it remains challenging to predict their microstructures. Using a near-atomistic, chemically accurate force field, we studied the phase behavior of chromatin regulators that are crucial for heterochromatin organization and their interactions with DNA. Our computed phase diagrams recapitulated previous experimental findings on different proteins. They revealed a strong dependence of condensate stability on the protein-DNA mixing ratio as a result of balancing protein-protein interactions and charge neutralization. Notably, a layered organization was observed in condensates formed by mixing HP1, histone H1, and DNA. This layered organization may be of biological relevance as it enables cooperative DNA packaging between the two chromatin regulators: histone H1 softens the DNA to facilitate the compaction induced by HP1 droplets. Our study supports near atomistic models as a valuable tool for characterizing the structure and stability of biological condensates.

## Introduction

Compartmentalization allows eukaryotic cells to coordinate biochemical reactions with high specificity and efficiency. In addition to traditional membrane-enclosed compartments, recent studies have uncovered a variety of membraneless organelles.^1–4^ These organelles are dynamic and can adjust their composition in response to ligand binding and chemical modifications,^5^ rendering them ideal designs for signal sensing. ^6^ Membraneless organelles are prevalent in the cytosol, with examples including P granules, ^7^ stress granules,^8^ etc. They are frequently found in the nucleus as well^9^ and may underlie the organization of heterochromatin,^10–12^ superenhancers, ^13^ nuclear speckles,,^14^ and nucleoli.^15,16^ Despite their distinct composition and cellular localization, membraneless organelles appear to share a common formation mechanism known as liquid-liquid phase separation (LLPS). ^17,18^ LLPS can be driven by multivalent interactions between intrinsically disordered proteins (IDPs), which are frequently found in the organelles.^19,20^

Besides IDPs, membraneless organelles often contain RNA and DNA molecules as well.^17,18,21^ The complex phase behavior of such multi-component systems is becoming increasingly appreciated.^22,23^ The presence of nucleotides can give rise to a scaffold-client mechanism to drive the phase separation of IDPs that do not aggregate on their own.^24,25^ The stability of the formed condensates exhibits nonmonotonic dependence on nucleotide concentration, giving rise to the so-called reentrant phase separation. ^26,27^ Further, multiple component phase separation supports novel outcomes not present in binary polymer-solvent mixtures. For example, the condensate composition and the molar fraction of individual components can be fine-tuned by adjusting their interaction strength.^23,28^ Rather than a homogeneous mixture, droplets may instead exhibit a layered organization with subcompartments, which has indeed been observed in nucleoli^15^ and speckles.^29^ Finally, multiple demixed droplets without interfacial contacts can also form. ^30–32^

General principles of multi-component phase separation are beginning to emerge. ^22,23^ Electrostatic interactions between charged molecules have received great attention, and the complex coacervation mechanism serves as a powerful theoretical framework for studying condensate stability. ^33,34^ Jacobs and Frenkel showed that demixed states could form in the presence of large fluctuations of intermolecular interaction strengths, facilitating the coexistence of multiple organelles.^30^ In addition to demixed states, Pappu and coworkers further found that single condensates but with layered organization and subcompartments could form for certain interaction patterns. ^15,35,36^ The precise morphology of the droplet with multiple coexisting phases is dependent on their interfacial tension and volume fractions.^37^ Despite these advances in understanding multi-component phase separation, predicting the microstructures of the multi-component condensates remains challenging. The variety of possible outcomes in spatial organization places a high demand on the model accuracy.

In this work, we simulated LLPS and the organization of multi-component systems with a near-atomistic model. We focused on the phase behavior of chromatin regulators that play crucial roles in heterochromatin formation and gene silencing, including heterochromatin protein 1 (HP1) and histone H1.^2,38^ A recently developed force field, MOFF,^39^ that is well suited for simulating both ordered and disordered proteins was combined with a chemically accurate DNA model^40^ to study protein-DNA condensates. Our simulations resolved the difference of various chromatin regulators in their phase behaviors and their association with DNA molecules. A linear model that accounts for both protein-protein interactions and system net charges can quantitatively reproduce the critical temperature for various systems and provides insight into the stability of biological condensates. Furthermore, DNA molecules can act as bridging agents to support layered organization in ternary systems that mix them with HP1*α* and H1. Such a layered organization promotes collaboration between the two chromatin regulators to compact DNA and possibly package heterochromatin. Our study on HP1 homologs further supports a model proposed by Keenen et al., ^41^ in which the formation of heterodimers may destabilize HP1*α*/DNA condensates. We conclude that near-atomistic modeling is useful for predicting the microstructures and phase behavior of multi-component systems and uncovering the design principles of biomolecular condensates.

## Results

### Near-atomistic modeling resolves the distinct phase behavior of chromatin regulators

To study multi-component LLPS of biological significance, we characterized the phase behavior of three chromatin regulator proteins, HP1*α*, HP1*β*, and histone H1 (Figure 1 and Table S1-S2), with DNA. All proteins consist of both folded and disordered regions. The two folded domains of HP1 include a chromodomain (CD) that recognizes methylated N-terminal tails of histone H3 (H3K9me3) and a chromoshadow domain (CSD) that enables dimerization.^42–44^ The globular domain of H1 was frequently found to bind near nucleosome dyad and contact linker DNA.^38,45^ Meanwhile, the disordered regions in these proteins can drive their LLPS when mixed with DNA.^24,41,46,47^ As such, they serve as ideal model systems to investigate how interactions between proteins and nucleic acids contribute to the stability of biological condensates.

**Figure 1:**
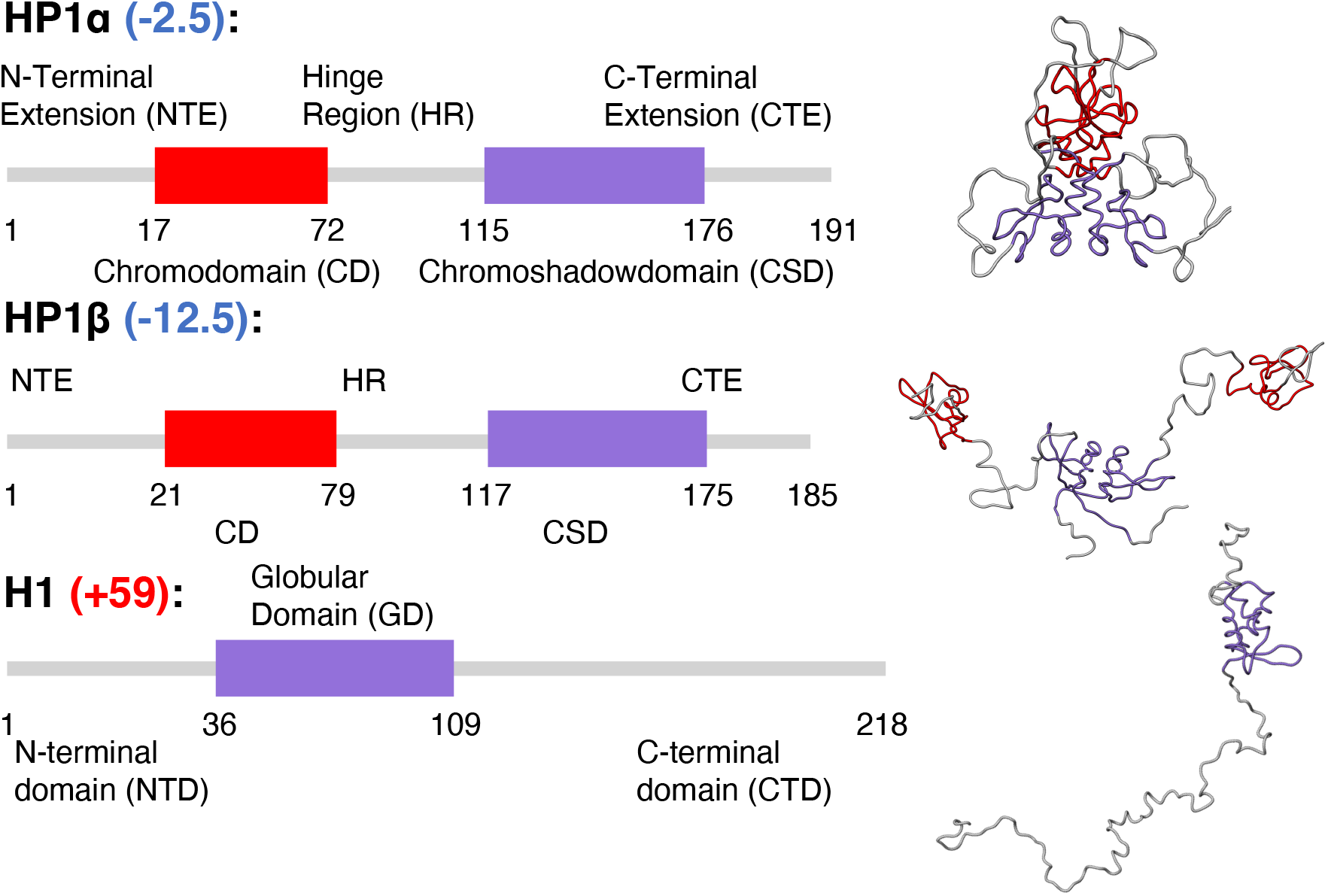
Overview of sequence and structural features of the three chromatin regulators. The boxes indicate folded domains, while grey lines correspond to disordered regions. Numbers in parentheses represent the net charge of each protein. The structures shown on the right are the most likely conformations determined from simulations^39^ and include dimers for HP1 proteins and a monomer for H1.

Structural characterization of chromatin regulators is challenging, and their inclusion of both folded and disordered domains places a high demand on the force field accuracy. We previously introduced MOFF,^39^ a coarse-grained protein force field that succeeds in predicting the size of disordered proteins and in folding certain globular proteins. When applied to HP1*α* and HP1*β*, MOFF predicted the radius of gyration (*R*_*g*_) with values that match small-angle X-ray scattering (SAXS) results. Further, consistent with the observation that HP1*α* can phase separate more readily than HP1*β*,^10,12,48,49^ the critical temperature of phase separation (*T*_*C*_) for HP1*α* determined using MOFF is higher than that for HP1*β*. We carried out slab simulations for histone H1 and found that it does not phase separate under all conditions we tested (Figure 2B and Figure S1). This result is in agreement with previous experiments,^46,50^ and is likely due to H1’s large net positive charge.

**Figure 2:**
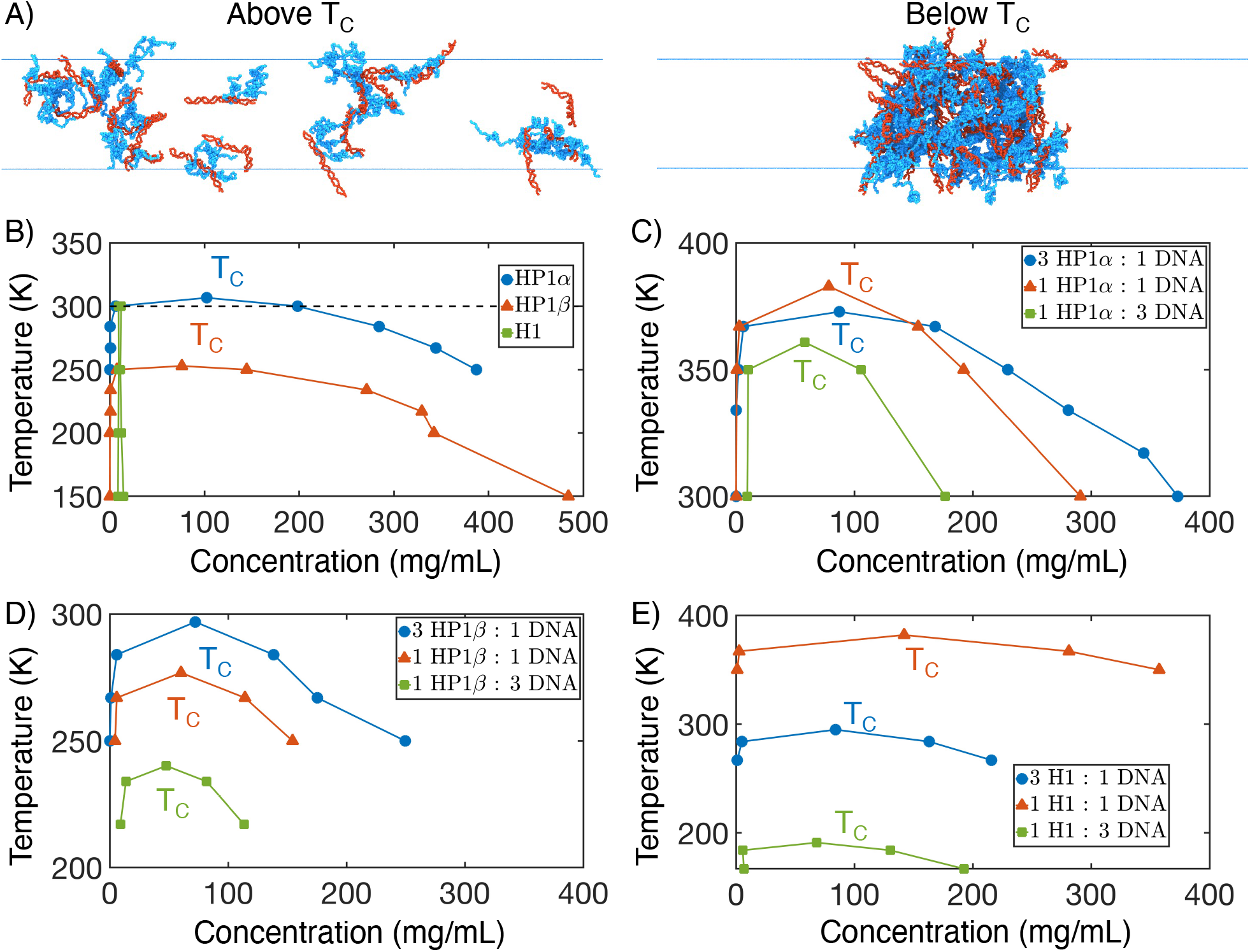
Phase diagrams of proteins and protein-DNA mixtures. (A) Example configurations of 1 HP1*α* (blue) : 1 DNA (red) above and below *T*_*C*_. (B) Phase diagrams of protein systems without DNA. The small concentration values for H1 indicate that it does not phase separate on its own. Results for HP1 homologs are adapted with permission from Ref. ^39^ Copyright 2021 American Chemical Society. (C-E) Phase diagrams and critical temperatures of HP1*α* (C), HP1*β* (D), and H1 (E) at different mixing ratios with DNA.

We further combined MOFF with the MRG-CG DNA model^40^ to study the impact of DNA molecules on protein phase separation. Accounting for the DNA is crucial for a more biologically relevant scenario since chromatin regulators primarily reside inside the nucleus filled with DNA. Numerous studies have shown that DNA could significantly alter both the thermodynamics and kinetics of phase separation.^51–53^ The DNA model represents each nucleotide with one bead and reproduces the persistence length at the physiological salt concentration (Figure S2). Protein-DNA interactions were adjusted to reproduce the binding free energy for a set of complexes (Figure S3). With this calibrated force field and the slab simulation methodology,^39,54^ we constructed phase diagrams for each protein at different ratios of protein (dimers of HP1 or monomers of H1) to a 50-bp-long DNA, while keeping the total number of molecules fixed at 100.

The three proteins exhibit striking differences in their interaction with DNA molecules (Figure 2). In agreement with experimental findings, HP1*α* and H1, but not HP1*β*, can phase separate at room temperature at certain protein-DNA ratios. ^10,24,41^ Notably, the response of *T*_*C*_ to DNA concentration varies dramatically from protein to protein. HP1*α* favors an equal protein to DNA ratio (1HP1*α*:1DNA), but these effects are relatively subtle, and the *T*_*C*_ stays between 361 K and 383 K for all mixing ratios. On the other hand, while H1 similarly favors the equal mixing ratio, its critical temperature changed from 382 K at 1H1:1DNA to 191 K at 1H1:3DNA. Indeed, experiments have shown that similar ratios are favored for both HP1*α* and H1. ^24,41^ Meanwhile, 3HP1*β*:1DNA is preferred over other setups.

### Sequence-specific interactions and charge balance dictate condensate stability

The apparent disparity in the protein-DNA ratio of the most stable condensates motivated us to search for a better theoretical understanding of their stability. One possible quantity of importance is the net system charge, *Q*. Through a complex coacervation mechanism, ^33,55^ for which electrostatic interactions between oppositely charged polyelectrolytes dominate the system, low levels of DNA can enhance condensate formation, whereas high levels can cause their dissolution. The optimal DNA ratio for maximum stability is expected to neutralize the overall charge of the condensate.^26–28^ While it may explain the results for H1, the charge-balance mechanism fails to reconcile the differences across proteins. In particular, HP1 molecules are negatively charged themselves and cannot approach a neutral droplet (Figure 3A), but nevertheless form more stable condensates with DNA than H1. As such, more refined mechanisms must be utilized to understand the properties of these condensates. One key component ignored in the above charge-balance argument is the short-range van der Walls interactions between protein molecules. From the Flory–Huggins theory of mixing for polymer solutions, ^56^ it is well known that *T*_*C*_ depends linearly on the interaction energy between proteins, *E*_pp_. Therefore, we included both factors in a linear regression model to fit *T*_*C*_ estimated from slab simulations and obtained

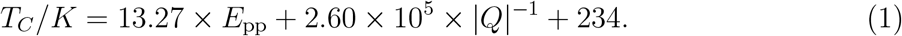

**Figure 3:**
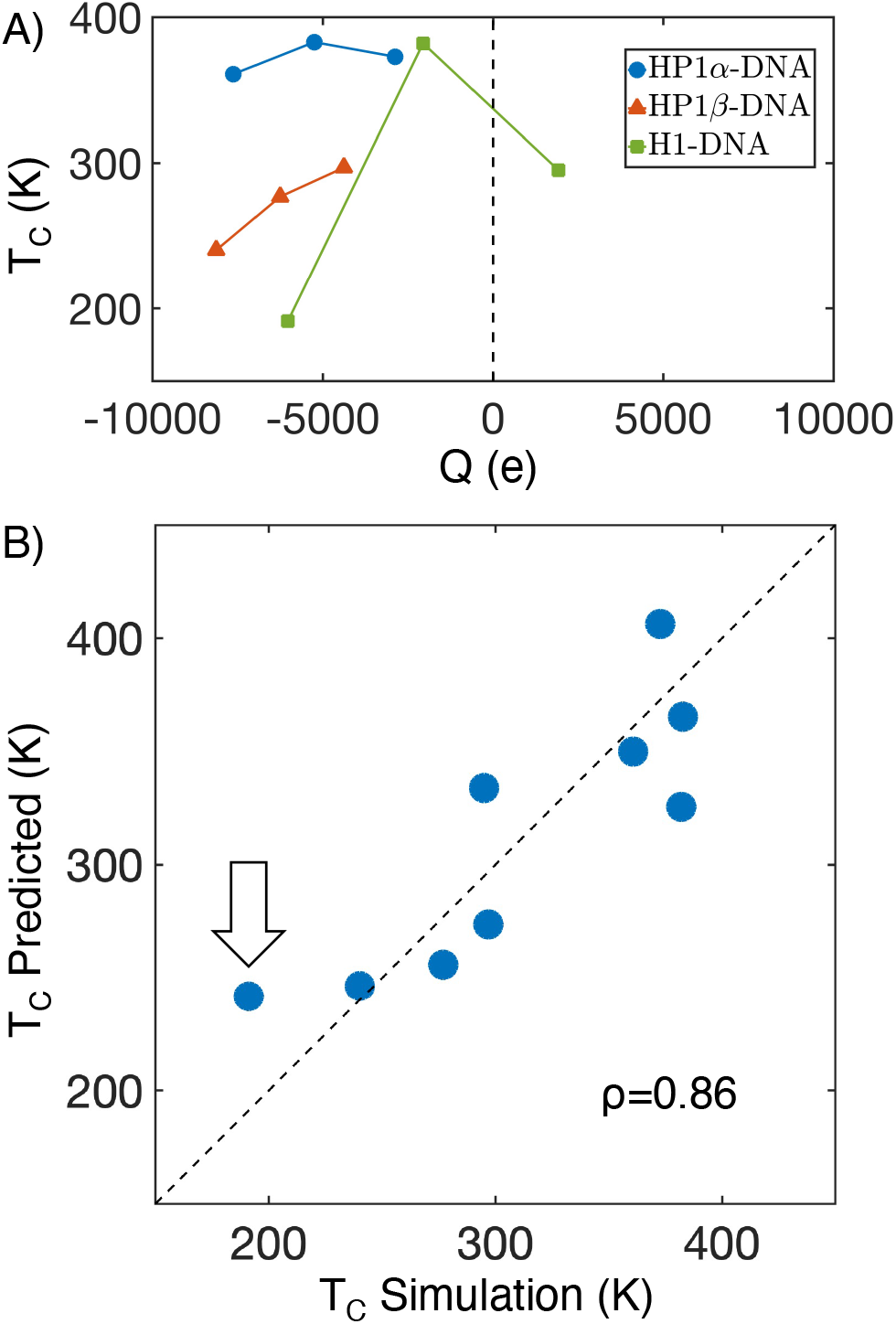
Predicting the critical temperature, *T*_*C*_, of multi-component phase separation. (A) *T*_*C*_ as a function of the net system charge (Q) for the three chromatin regulators mixed at different ratios with DNA molecules. (B) Comparison between simulated *T*_*C*_ and values obtained through linear regression, given in Eq. 1. *ρ* is the Pearson correlation coefficient between the two data sets, and the dashed diagonal line was provided as a guide to the eye. The arrow draws attention to the largest outlier, 1H1:3DNA.

We estimated *E*_pp_ from the binding free energy curves for two HP1 dimers or two H1 monomers (Figure S4).

This linear model for *T*_*C*_ fits our data well (Figure 3B), while neither *Q* nor *E*_pp_ alone can provide fits of comparable quality (Figure S5). Further, it provides an intuitive explanation of our simulated trends. For example, HP1*α*-DNA condensates are stable across many protein-DNA ratios due to the strong protein-protein interactions and HP1*α*’s relatively small net charge. In comparison, condensates formed with HP1*β* are less stable as the corresponding *E*_pp_ is negative, and the protein is more negatively charged. Finally, the strong dependence of H1-DNA condensate stability on the protein-DNA ratio arises from the competition between the repulsive H1-H1 and attractive protein-DNA interactions.

However, given its simplicity, we do not expect the linear model to account for all the complex behaviors of protein-DNA condensates. In particular, a mean-field model of the electrostatic contribution to the free energy based on charge density is insufficient for polyelectrolytes. ^57^ The polymer topology places significant constraints on charge distribution.

Both HP1*α* and HP1*β* bind favorably with DNA despite their overall negative charge (Figure S6). Therefore, the simple dependence on net charge cannot explain the enhanced phase separation upon introducing DNA because of favorable protein-DNA interactions. In addition, the tight binding between histone H1 and DNA may give rise to magic number effects.^58,59^ Contributions from configurational entropy to phase separation in such systems become essential but are not captured by either *E*_pp_ or *Q*. Therefore, H1 systems often appear as outliers of the linear fit.

### DNA bridging drives layered organization in ternary systems

As they coexist inside the nucleus, chromatin regulators may cooperate or compete when binding with DNA. Prior *in vivo* studies have suggested that HP1 and H1 localize together in heterochromatic regions.^50^ The microscopic structures of such ternary systems are poorly characterized. Recent studies have shown that three-component systems with solvent can give rise to complex multiphase scenarios, including demixed droplets, a condensed homogenous phase, or a layered organization with sub-compartments. ^35,60^ We determined the temperature-density phase diagrams for ternary complexes to study both their structural organization and stability.

We performed slab simulations with 1HP1*α*:1H1:2DNA and 1HP1*α*:1HP1*β*:2DNA at various temperatures to probe the density of condensed and dilute phases (see Table S1 for simulation details). As shown in Figure 4, both systems display upper critical temperatures comparable to 1HP1*α*:1DNA. Therefore, H1 and HP1*β* do not seem to impact the stability of condensates formed by HP1*α* and DNA. This observation is in strong contrast to the phase diagrams of protein-only mixtures. As shown in Figure S7, both H1 and HP1*β* destabilize condensates formed only with HP1*α*, preventing LLPS at room temperature. The destabilizing effect likely stems from the repulsive interactions between H1-H1 and HP1*β*-HP1*β* (Figure S4). The shape of the ternary complex phase diagrams also appears different from that of protein-DNA condensates shown in Figure 2. For example, two regimes can often be seen. The two regimes are particularly visible for 1HP1*α*:1HP1*β*:2DNA, supporting the definition of two critical temperatures (Figure S8).

**Figure 4:**
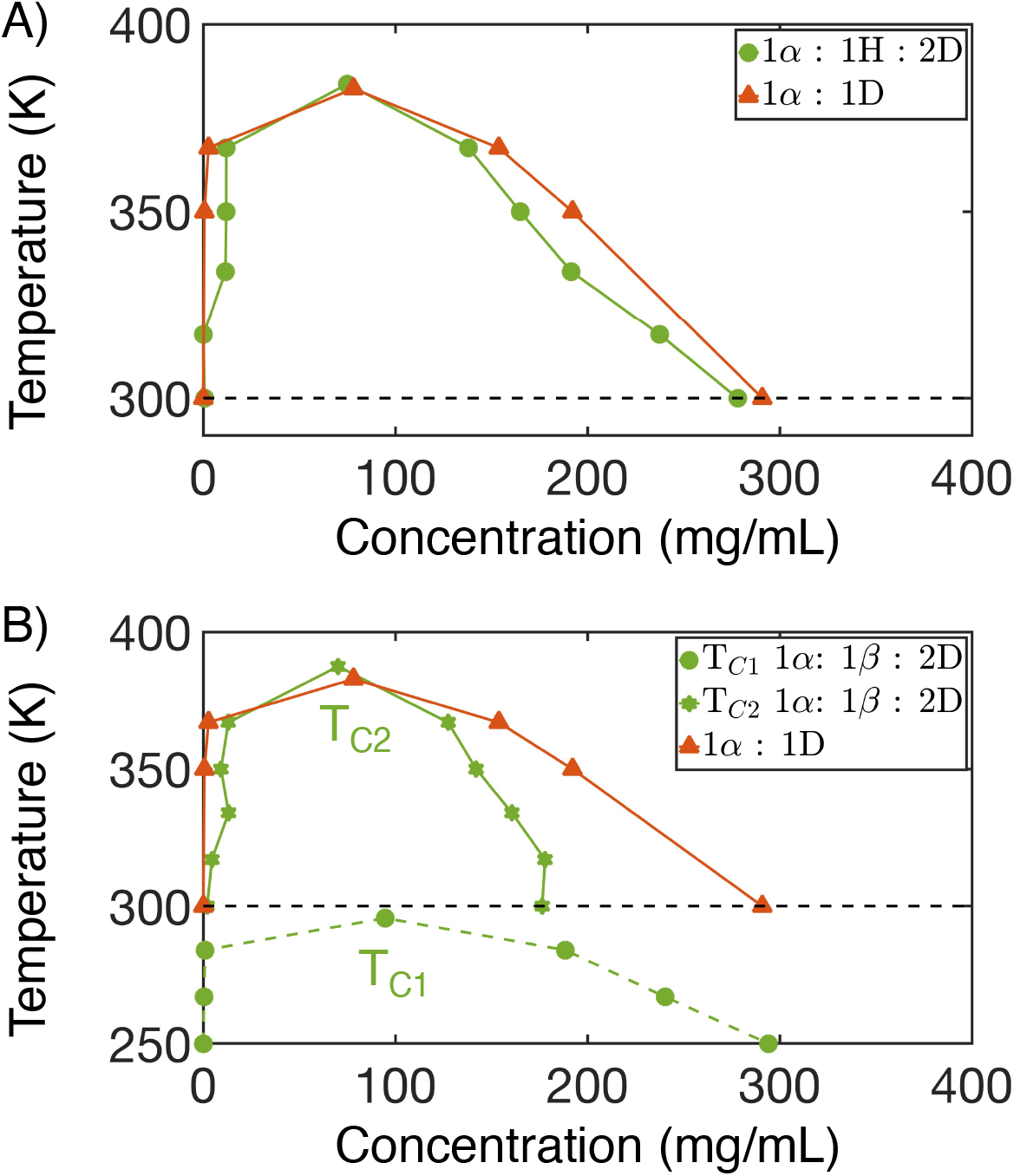
Phase diagrams of ternary systems with implicit solvation for HP1*α*-H1-DNA (A) and HP1*α*-HP1*β*-DNA (B) mixtures. Abbreviations are used for different biomolecules (*α* - HP1*α*, H - H1, D - DNA, *β* - HP1*β*). The result for 1HP1*α*:1DNA is included for comparison.

We further examined the structures to better understand the stability of the ternary mixtures and interpret the phase diagrams. At low temperatures, both systems formed a layered organization, which corresponds to a demixed state with two contacting liquid phases that differ in their relative composition.^35^ The layered organization can be readily seen in the *z*-density profile of the simulated condensates. For both systems, HP1*α* is concentrated in the middle of the slab, with H1 or HP1*β* residing at the exterior (Figure 5A,C). The inclusion of DNA in both liquids reduces the surface tension between them to prevent complete separation. The stability of the two systems differs dramatically, however. For example, the condensate with H1 remains stable at 300 K, supporting the biological relevance of the layered organization. On the other hand, the condensate with HP1*β* tends to shed off HP1*β* at high temperatures, resulting in more homogeneous phases that are increasingly enriched with HP1*α* (Figure 5D). We note that, because of the strong electrostatic interactions in these systems, simulations starting from different initial configurations produced slightly different *z*-density profiles. However, the formation of layered organizations is robust and can be seen in all simulations (Figure S9).

**Figure 5:**
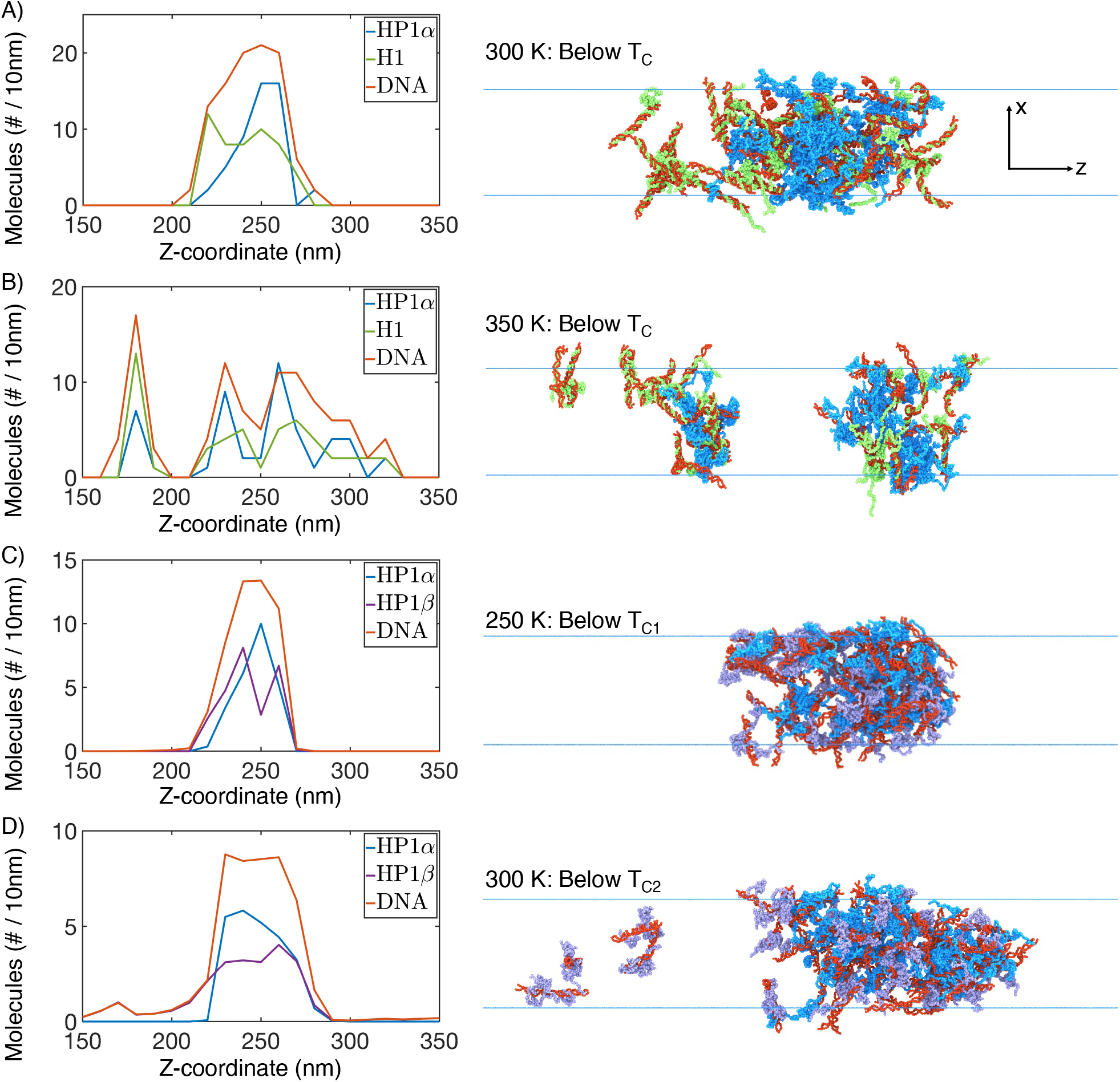
Slab density profiles support layered organizations in ternary mixtures. (A) At 300 K for 1HP1*α* (blue) : 1H1 (green) : 2DNA (red), HP1*α* coalesces toward the center of the droplet, with H1 to the outside. (B) Still below *T*_*C*_ for 1HP1*α*:1H1:2DNA at 350 K, the layered organization becomes less evident and molecules are more uniformly distributed in the condensates. (C) Below *T*_*C*1_ (250 K) for 1HP1*α* (blue) : 1HP1*β* (purple) : 2DNA (red), HP1*α* coalesces toward the center of the droplet, with HP1*β* to the outside. (D) Below *T*_*C*2_ (300 K) for 1HP1*α*:1HP1*β*:2DNA, HP1*β*/DNA complexes begin to fall off the condensate, leaving behind a state enriched in HP1*α*.

Therefore, the condensates of ternary systems are made up of two liquid phases enriched and depleted of HP1*α*. Their temperature-density phase diagrams can indeed be understood from studies on partially miscible liquids.^61^ For HP1*α*-HP1*β*-2DNA, the system first undergoes a “boiling” process to dissociate HP1*β*/DNA complexes at *T*_*C*1_ (see Figure 4B and Figure 5D). The second transition corresponds to the vapor-liquid phase separation of HP1*α* and DNA, and the corresponding *T*_*C*2_ is comparable to that of the binary mixtures. On the other hand, the two phases in HP1-H1-2DNA become more miscible at higher temperatures before the whole condensate falls apart, and the boiling transition is less evident. The difference between the two systems partially arises from the more favorable interaction between HP1*α* and H1 than that between HP1*α* and HP1*β* (Figure S10).

### Layered organization facilitates DNA softening and compaction

While slab simulations combined with phase diagrams can highlight the diverse interactions that drive phase separation, they provide little information regarding the impact of liquid droplets on chromatin conformation due to our use of short DNA segments. Cellular environments likely mirror a single long DNA strand, the conformation of which may depend strongly on the contacting chromatin regulators. ^40^ To examine whether the condensates studied here can compact chromatin to restrict DNA accessibility, we performed simulations for a 200-bp-long DNA with the presence of protein droplets. While chromatin differs from naked DNA due to its incorporation of histone proteins, the simulations may nevertheless provide insight into the impact of polymer topology on phase separation and vice versa.

Starting from a conformation with 100 HP1 dimers positioned next to the DNA molecule, we performed long-timescale simulations to equilibrate the system (Figure 6A). We found that HP1*α* condensed onto the DNA (Figure S11A), resulting in a large droplet that incorporates most proteins (Figure 6C). The protein droplet induced highly bent DNA configurations (Figure 6A), giving rise to a second maximum at smaller values in the distribution of the DNA end-to-end distance (Figure S11). Notably, similarly collapsed DNA configurations do not appear metastable upon binding of one or two HP1*α* dimers (Figure S12), indicating the importance of a collective role by many proteins in bending the DNA. DNA bending by HP1*α* droplets has indeed been observed by Keenen et al. using single-molecule DNA curtain experiments. ^41^ However, a quantitative comparison between simulation and experimental results may be challenging since the absolute stability of the collapsed DNA configuration depends on both the DNA length and the protein concentration.

**Figure 6:**
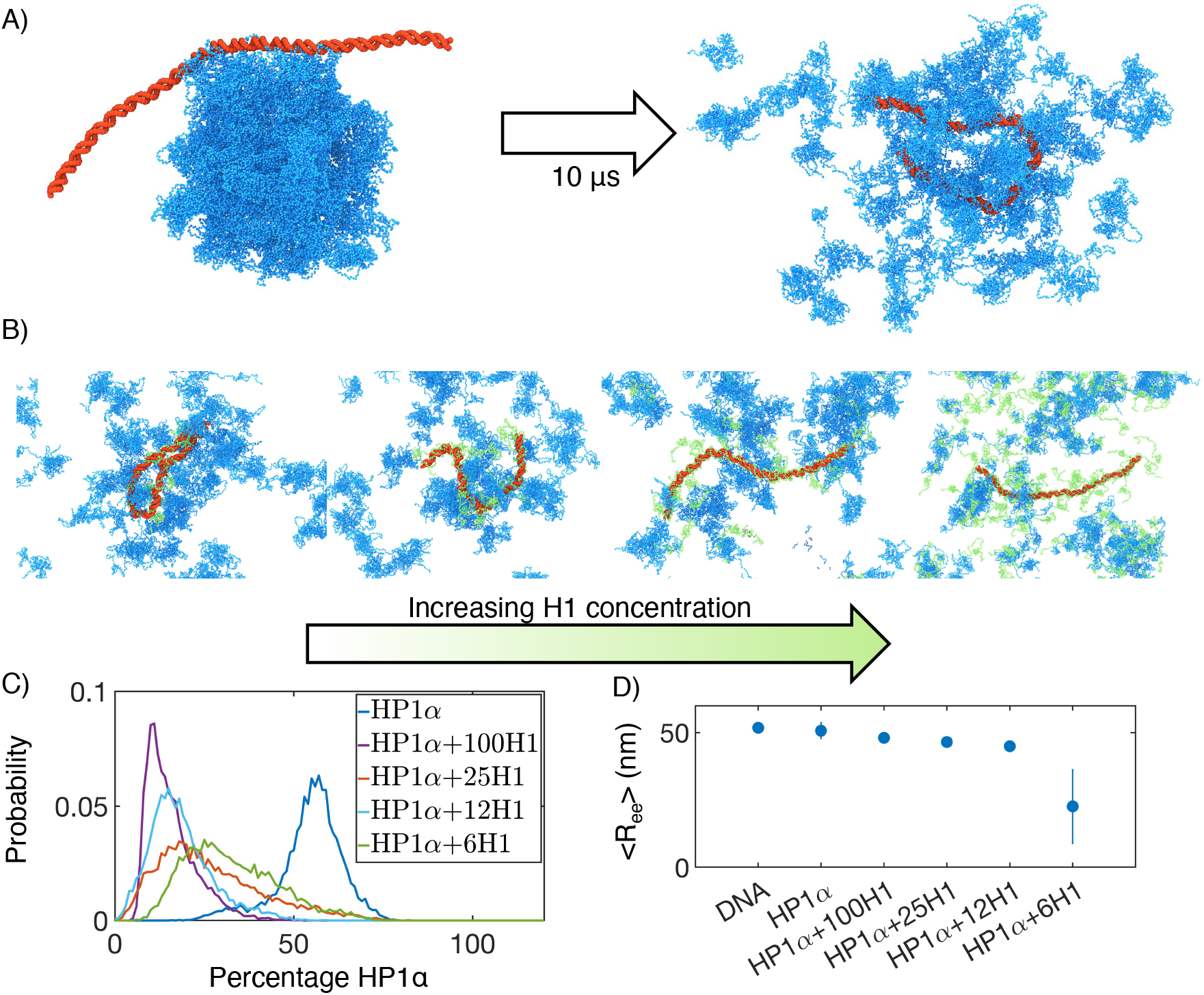
Impact of protein droplets on single DNA conformation. (A) Illustration of the simulation protocol for studying the impact of protein (blue) droplets on the conformation of a 200 bp long DNA (red). (B) Snapshots from simulations of HP1*α*+H1+DNA mixtures, with the H1 (green) concentration increasing from left to right. (C) Impact of H1 concentration on HP1*α*’s participation in cluster formation. Different curves represent probability distributions of the percentage of HP1*α* in the system that is incorporated into the largest cluster formed by protein and DNA molecules. (D) Impact of LLPS on the average DNA end to end distance (⟨*R*_ee_⟩). A simulated result under identical conditions but without proteins is also included for comparison (DNA). Error bars represent standard deviations of the mean.

To examine how HP1*α*-H1 mixtures might affect DNA conformations, we performed simulations with varying additional amounts of H1: from 100 to 6 monomers. Consistent with those presented in Figure 5, the single-DNA simulations also resulted in layered organizations, with DNA-bound H1 surrounding HP1 condensates (Figure 6B). However, due to their stronger binding affinity (Figure S6), H1 molecules essentially cover the entire DNA. At high concentrations, H1 can even strip HP1*α* away from the DNA (Figure S11A), reducing HP1*α*’s participation in droplet formation (Figure 6C).

In addition to forming layered organizations, we noticed that introducing H1 induced more DNA bending. A smaller average end-to-end distance is particularly evident at lower H1 concentrations. By coating the DNA, H1 molecules effectively screen the repulsive interactions between nucleotides with their positive charges (Figure 6D, Figure S11B).^62^ Furthermore, at lower H1 concentrations, HP1*α* remain bound to the DNA to stabilize highly bent conformations through phase separation (Figure 6B, far left). Therefore, HP1*α* and H1 can work collaboratively to control and condense DNA.

## Discussion

We utilized two recently developed near-atomistic force fields^39,40^ to study the stability and structural organization of protein-DNA condensates. The simulations succeeded at resolving the difference in phase behaviors of three chromatin regulators and qualitatively reproducing experimental observations. We discovered that the stability of various condensates could be quantitatively captured with a surprisingly simple, linear model that accounts for protein-protein binding free energy and system net charge. In addition, a ternary system that includes HP1*α*, H1, and DNA with implicit solvation phase separates at room temperature into condensates with a layered organization. This layered organization arises due to the bridging of two phases enriched with H1 and HP1 respectively by DNA molecules. Finally, we observed that protein condensates could affect DNA conformation, with H1 molecules softening the DNA while HP1*α* stabilizing compact structures.

Our results have significant implications on heterochromatin organization *in vivo*. The layered droplets formed by HP1*α*, H1, and DNA could persist with the presence of chromatin. In particular, the higher DNA affinity of H1 molecules could allow them to localize to the interior of the chromatin fiber, where H1 binds near nucleosome dyads and with linker DNA.^45^ Meanwhile, HP1*α* would favor configurations closer to the exterior of the fiber, where it could still recognize H3K9me3 marks. ^63^ Such a layered distribution could facilitate the two molecules to collaboratively soften and compact chromatin. While our focus in this study is on protein-DNA droplets, we hope to extend these models to chromatin structures in the future.

### HP1*β* destabilizes HP1*α*-DNA droplets through heterodimerization

We note that the phase diagram of the ternary system, 1HP1*α*:1HP1*β*:2DNA, is not entirely consistent with observations made by Keenen et al. ^41^ The authors found that HP1*β* can dissolve HP1*α*-DNA droplets, suggesting less stable condensates and a lower *T*_*C*_ for the ternary system compared to HP1*α*-DNA. However, the phase diagrams showed that the two systems share comparable values for *T*_*C*_. The discrepancy could arise from the inability of our simulations to capture heterodimerization that might occur on a much slower timescale after the homodimers dissociate. As suggested by Keenen et al., ^41^ heterodimerization could break the interactions that stabilize HP1*α* droplets. To test this hypothesis while avoiding simulating the slow kinetics of heterodimer formation from homodimers, we performed additional simulations in which the HP1 homologs were restricted in heterodimeric conformations. We found that the radius of gyration of the heterodimer lies in between that of HP1*α* and HP1*β* (Figure S13).^39^

We computed the phase diagram and *T*_*C*_ for pure heterodimers, which we denote as HP1*α*/HP1*β*, and their DNA mixture (Figure S14). While the condensate formed by heterodimers is equally stable compared to that of HP1*α* and HP1*β* homodimers, the stability of the DNA mixture is significantly reduced. However, we still predicted LLPS at room temperature for 1HP1*α*/HP1*β*:1DNA. While this could be due to the limited resolution of our coarse-grained model, a more likely possibility is the use of a high DNA concentration in our slab simulations. We next turned to simulations with a long, single DNA (Figure S15), where the protein-DNA ratio is more comparable to the experimental value. ^41^ In these simulations, the heterodimer significantly reduced the percentage of HP1*α* in the largest cluster formed by protein and DNA molecules relative to homodimer simulations (HP1*α*+HP1*β*). Therefore, our simulations indeed support heterodimerization as a mechanism to destabilize HP1*α* droplets formed with DNA.

## Methods

### Near-atomistic protein-DNA force fields

Simulating large-scale phenomena such as LLPS necessitates balancing computational expense with model resolution. We represented protein and DNA molecules with one coarse-grained bead per amino acid or nucleotide. Models of similar resolutions have been widely used to study IDPs^64–68^ and protein-DNA complexes^69–77^ with great success. The MOFF force field for proteins^39^ and molecular renormalization group coarse-graining model (MRG-CG) for the DNA^40^ were combined together to describe protein-DNA interactions under implicit solvation. Each model is described briefly below, with full details provided in the Supporting Information (SI). An implementation of our force field can be found at the [group GitHub page](https://github.com/ZhangGroup-MITChemistry/MOFF/tree/main/MOFF_DNA).

The potential energy for a protein structure ***r*** in MOFF is defined as

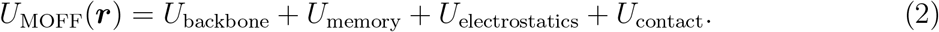

The first two terms are responsible for maintaining the backbone geometry and secondary structures of protein molecules. Equilibrium distances and angles used in defining these potentials were determined from initial structures built with RaptorX. ^78^ *U*_electrostatics_ describes electrostatic interactions between charged residues with the Debye-Hückle theory and a distance-dependent dielectric constant. The last term *U*_contact_ is the amino acid type-dependent pairwise contact potential optimized to reproduce the radius of gyration for a list of folded and disordered proteins. In addition, for proteins with folded domains, structure-based potentials^79^ were introduced to stabilize tertiary contacts. The strength of these potentials was tuned to reproduce the root mean squared fluctuations estimated with all-atom simulations (Figure S1).

The potential energy of the MRG-CG DNA model takes the form

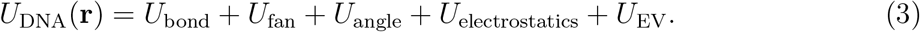

*U*_bond_, *U*_fan_, and *U*_angle_ account for local interactions among nearby nucleotides, while *U*_electrostatics_ and *U*_EV_ correspond to nonbonded, electrostatic interactions and the excluded volume effect. While the original model was parameterized with explicit ions, we switched to implicit ions to be consistent with the electrostatic potential defined in the protein force field. After adjusting the strength of local interactions, we found that the implicit ion model succeeded in predicting the DNA persistence length at the physiological salt concentration (Figure S2).

Protein-DNA interactions included electrostatic and excluded volume contributions. This treatment worked well at reproducing the experimental binding free energies for various protein-DNA complexes (Figure S3).

### Molecular dynamics simulation details

We carried out molecular dynamics simulations using the software GROMACS^80^ with a timestep of 10 fs.

We used the slab simulation technique to probe the thermodynamic stability of condensates and determine the critical temperature *T*_*C*_.^39,54,65^ Via this technique, interfaces of a dense biopolymer phase and a dilute vapor (implicit solvent) phase were simulated at various possible temperatures to determine temperature-density phase diagrams. Each simulation lasted for two *µs* and Langevin dynamics was used to control the temperature. We fixed the total number of molecules, including HP1 dimers, H1 monomers, and dsDNA, to be 100 for all slab simulations at different protein-DNA mixing ratios. The dsDNA used in these simulations is 50 bp long, a length that is short enough to avoid interactions with its periodic image but long enough to bind H1 and HP1. The exact protein-DNA ratios and temperatures of our simulations are provided in Table S1. Additional simulation details can be found in the *SI Section: Computing phase diagrams with slab simulations*.

In addition to slab simulations, we studied the effect of protein phase separation on DNA conformation. These simulations began with a strand of 200-bp-long dsDNA bound to a droplet of 100 HP1*α* dimers inside a cubic box. We further introduced H1 or HP1*β* at various concentrations before carrying out ten-*µs*-long constant temperature simulations at 300 K. The first one *µ*s of these simulations were discarded for equilibration, leaving nine *µ*s of data to analyze. These simulations are described in full in the *SI Section: Probing the impact of phase separation on DNA conformation*.

## Acknowledgement

This work was supported by the National Institutes of Health (Grant 1R35GM133580) and the National Science Foundation (Grant MCB-2042362). A.L. further acknowledges support by the National Science Foundation Graduate Research Fellowship Program. We thank Dr. Xingcheng Lin for his help implementing the MRG-CG DNA model.

